# ABC-finder: A containerized web server for the identification and topology prediction of ABC proteins

**DOI:** 10.1101/2020.11.18.042887

**Authors:** Poonam Vishwakarma, Naveen Kumar Meena, Rajendra Prasad, Andrew M. Lynn, Atanu Banerjee

## Abstract

In view of the multiple clinical and physiological implications of ABC transporter proteins, there is a considerable interest among researchers to characterize them functionally. However, such characterizations are based on the premise that ABC proteins are accurately identified in the proteome of an organism, and their topology is correctly predicted. With this objective, we have developed ABC-finder, i.e., a Docker-based package for the identification of ABC proteins in all organisms, and visualization of the topology of ABC proteins using a web browser. ABC-finder is built and deployed in a Linux container, making it scalable for many concurrent users on our servers and enabling users to download and run it locally. Overall, ABC-finder is a convenient, portable, and platform-independent tool for the identification and topology prediction of ABC proteins.

ABC-finder is accessible at http://abc-finder.osdd.jnu.ac.in

## INTRODUCTION

ABC proteins comprise one of the largest and most important protein families, most proteins being involved in active transport [1]. A typical ABC transporter is composed of a pair of nucleotide-binding domains (NBDs) and transmembrane domains (TMDs) [2]. While the NBDs are involved in fueling the transport process by means of ATP hydrolysis, the TMDs are responsible for substrate recognition and form the translocation channel [3]. NBDs have several conserved motifs, namely, Walker A, Walker B, Signature sequence (C-motif), H-loop, D-loop, and Q-loop [4]. Contrarily, the TMDs display poor conservation across different subfamilies. HUGO Gene Nomenclature Committee divided the ABC superfamily into 7 subfamilies from ABCA to ABCG. This classification was later extended for non-mammalian proteins with the addition of the ABCH subfamily found in insects [5] and fishes [6,7], and the ABCI subfamily found in plants [8]. ABC transporters can function both as importers as well as exporters, however, the former is restricted only to bacteria and plants and mediate the uptake of various nutrients, micronutrients, and phytohormones, etc. [9]. On the other hand, the ABC exporters facilitate the extrusion of a wide spectrum of substrates including ions, lipids, peptides, toxic xenobiotics, etc. [10]. Numerous studies have suggested the role of ABC transporters in various human diseases and chemoresistance [1]. Besides their role as transporters, some ABC proteins harbor just the NBDs, for instance, the ABCE and ABCF family representatives, and have implications in ribosome biogenesis, translation control, etc. [11,12], further adding to the immense biological relevance of this superfamily. The primary requirement for in-depth investigations pertaining to ABC proteins is their accurate identification from the genome/proteome and analysis of its topology. Even though programs performing certain steps in isolation exist, there is a need for a unified package to do the job with lesser hassles and more emphasis on reproducibility, keeping in view that reproducibility of research is the key element in modern science [13]. In an effort to make a simpler program available to the biology researcher community, we herein present “ABC-finder” as a simple and fast tool for identification and topology prediction of ABC proteins based on our previously established pipeline which led to the inventorization of ABC proteins in a number of yeast species [14–16]. ABC-finder combines stand-out methodologies, namely profile-HMM and TOPCONS for homology detection, and prediction of the topology of membrane proteins, respectively in a seamless manner whereby the users need to provide only the organism’s name or the proteome file as an input. Our analysis with reference organisms shows a strong correlation with the ABC protein inventories available in the literature.

## METHODOLOGIES AND WORKFLOW

### Data submission

Protein sequence data can be submitted in the following manner to the ABC-finder platform. The user can either specify the organism’s name (*Homo sapiens, Arabidopsis thaliana*, or *Candida glabrata*) or can upload a raw FASTA file (uncompressed) of the proteome. The file size is, however, limited to 95 MB. In the last step, the user needs to provide an email address to get the result files directly delivered.

### Method

#### Prediction of potential ABC proteins based on NBD HMM

To extract the putative ABC proteins from the sequence input, we utilized a protocol that was developed previously by us for inventorization of ABC proteins in yeasts [14–16]. Briefly, the HMM profile of the ABC-tran model (accession PF00005) obtained from the Pfam database [17] is used as a query to match against the proteome of an organism with the help of the “hmmsearch” function within the HMMER package [18] using the default settings. Positive hits above the default threshold are then further filtered based on a cutoff defined from the plot of domain-score and E-value. The default cut-off is system-generated from the E-value and requires manual filtering to remove proteins incorrectly classified above the cut-off. To automate this step, we use the observation that there is a small difference between the scores and E-values for proteins that belong to the same fold, but a large difference when scoring proteins from unrelated folds. This sudden drop in the sorted scores is exploited to mine proteins belonging to the defined class with minimal false positives.

#### Clustering of sequences using Cd-hit

Next, ABC-finder uses the Cluster Database at High Identity with Tolerance (Cd-hit) [19]. This clustering algorithm allows the grouping of all the sequences based on their sequence similarities. Given that Cd-hit clusters the highly similar (redundant) sequences, it allows the grouping of sequences resulting from different transcript variants of each ABC protein that usually populate the database in case of higher eukaryotes. The user also has the option to optimize the input parameters for the program from the “advanced options” menu.

#### Prediction of the TMD and protein topology

Since most of the families under the ABC superfamily include membrane transporters, it is relevant to detect their presence among putative proteins. Following the Cd-hit run, ABC-finder utilizes the stand-out program TOPCONS [20], which is a widely used program for consensus prediction of membrane protein topology. Once the TM helices are defined using TOPCONS, ABC-finder uses a set of in-house written Python scripts to demarcate the NBDs and TMDs in each of the predicted ABC proteins.

#### Visualization

E-value and domain score values of all the proteins above the inclusion threshold, as obtained from the HMM analysis are presented in the form of a plot. This plot allows the further filtering of sequences as defined in the “Prediction of potential ABC proteins based on NBD HMM” section. The arrangement of TMDs and NBDs as obtained with the help of TOPCONS is used to generate the final topology diagram of the proteins, respectively. Dynamic and Static plots are generated with Plotly (https://plot.ly/) and Orca (https://plot.ly/python/orca-management/). All the results files and several other log files are generated and zipped as the output data for downloading.

The overall workflow implemented in the ABC-finder server is shown in **Figure 1**. The help page on the web server provides a step-by-step guide to ABC-finder. All results are kept on the server for ten days.

**Figure 1:**
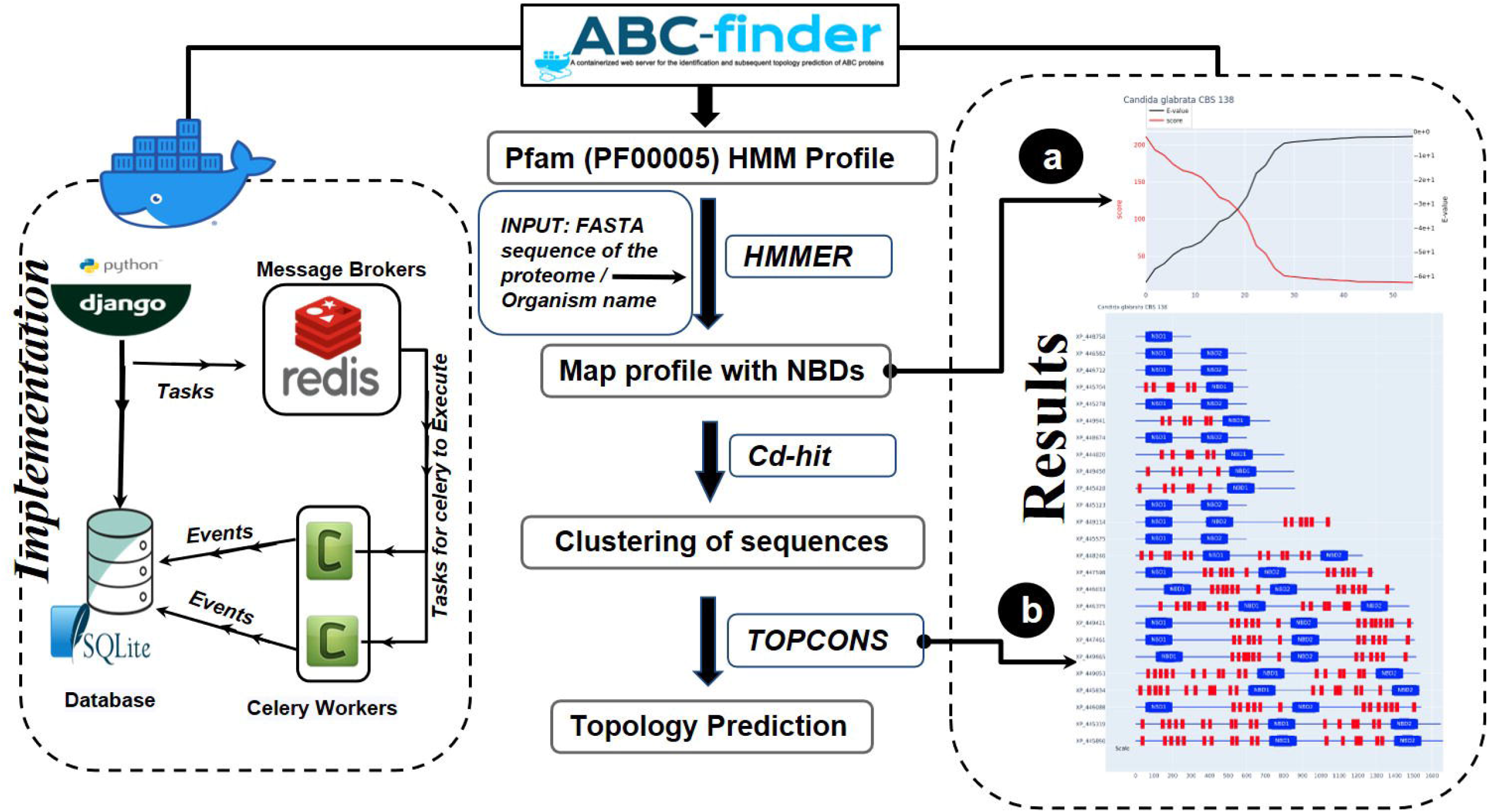
Generalized workflow of ABC-finder.

## IMPLEMENTATION

ABC-finder server is implemented in Python 3.7.6 (http://www.python.org/) using the Web framework Django 3.0.3 (http://www.djangoproject.com/). The system is containerized using Docker 19.03.3 [21] and docker-compose 1.25.0. Job queuing is carried out by the asynchronous task queue Celery 3.0.19 that uses the distributed message passing system Redis 3.3.11.

Users can also install the ABC-finder web server directly from the source code or build and run it from within Docker container using the instructions provided in the **Supplementary information**. The **Supplementary information** also contains FAQs and some useful links.

## RESULTS AND DISCUSSION

Once the input proteome file is uploaded or the organism is specified to ABC-finder, the server provides a zipped result folder that includes the following files in different formats (indicated within the parenthesis):

1. **search(.faa)** → it is the proteome file that is submitted to ABC-finder for analysis. It is either uploaded by the user or retrieved by the web server depending upon the organism specified by the user.
2. **profile(.hmm)** → it is the profile HMM derived from the PF00005 seed alignment.
3. **hmm_output** → it is the output obtained after running the “hmmsearch” function with the user-specified proteome file using profile.hmm as the query.
4. **E-value_Domain_Score(.html/.pdf/.PNG/.SVG/.JPEG)** → it is the plot of E-value and domain score for the sequences above the inclusion threshold as obtained from the hmm_output file. A representative plot for the analysis of *S. cerevisiae* proteome is provided in **Figure 2**.

**Figure 2:**
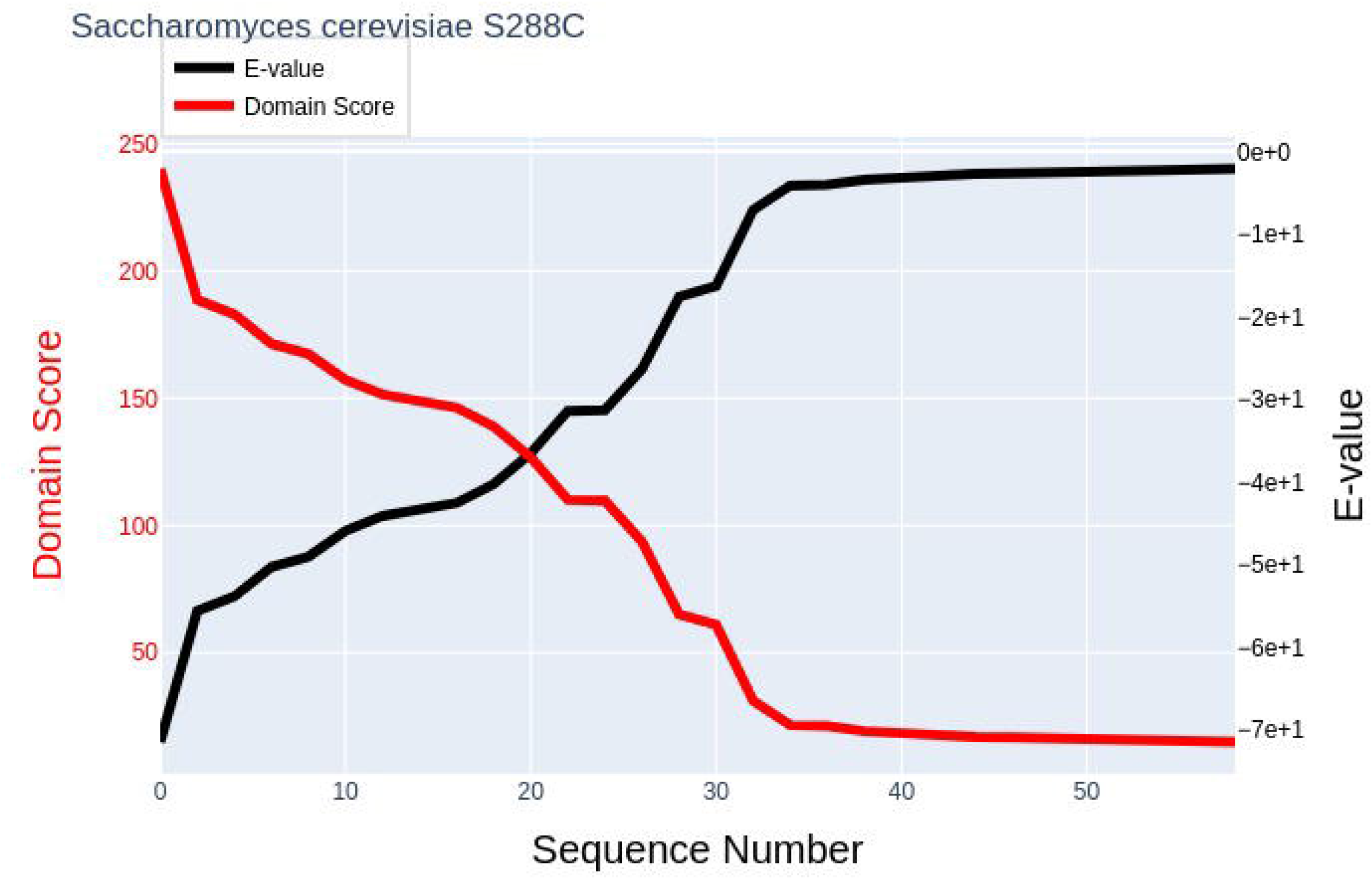
Plot of E-value and domain score of the sequences above the inclusion threshold value obtained after running the “hmmsearch” function with the *S. cerevisiae* S288C proteome.
5. **threshold(.txt)** → this file lists the IDs of the putative ABC protein candidates that are identified in the query proteome. These are predicted based on a cutoff defined from the plot of domain-score and E-value.
6. **search_faa0(.clstr)** → it is the output file obtained after running Cd-hit. It contains the different protein clusters based on sequence similarity. Users will need to refer to this file in special cases where multiple variants are reported for each protein in the database.
7. **search_faa1(.faa)** → FASTA file utilized as the input for TOPCONS.
8. **query.result(.txt)** → it is the final output file obtained from TOPCONS.
9. **Topology Plot(.html/.pdf/.PNG/.SVG/.JPEG)** → this plot represents the topology of the identified ABC proteins. TOPCONS is utilized for the prediction of the transmembrane domains. A representative topology plot for the predicted ABC proteins in the *S. cerevisiae* S288C strain is provided in **Figure 3**. The topology plot should be referred alongside the Cd-hit output file “search_faa0.clstr” to detect the highly similar sequence clusters, especially in the case of higher eukaryotes, wherein sequences from all the transcript variants of each protein are present in the database.

**Figure 3:**
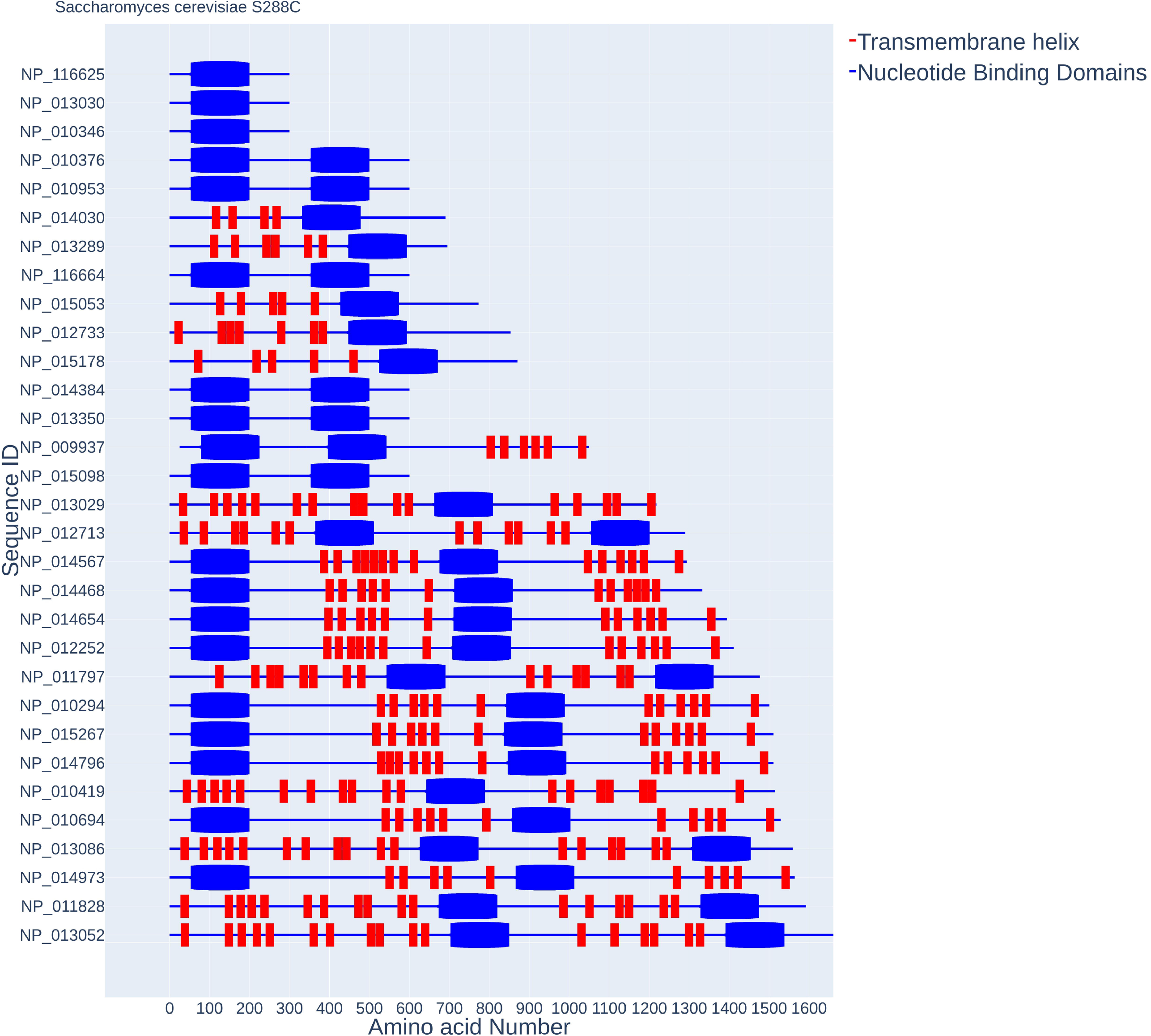
Plot representing the topology of the predicted ABC proteins in the *S. cerevisiae* S288C strain. While the x-axis denotes the amino acid number, the y-axis denotes the sequence IDs. TOPCONS is utilized for the prediction of the transmembrane domains. Red columns represent transmembrane domains and blue spheres represent the nucleotide binding domains.

To evaluate the performance of ABC-finder, we utilized already reported inventories of ABC proteins from prokaryotic as well as eukaryotic systems. The results are summarized in **Table 1**. Overall, the results show a very good correlation with the number of ABC proteins that are reported in the literature for the reference organisms.

**Table 1:**
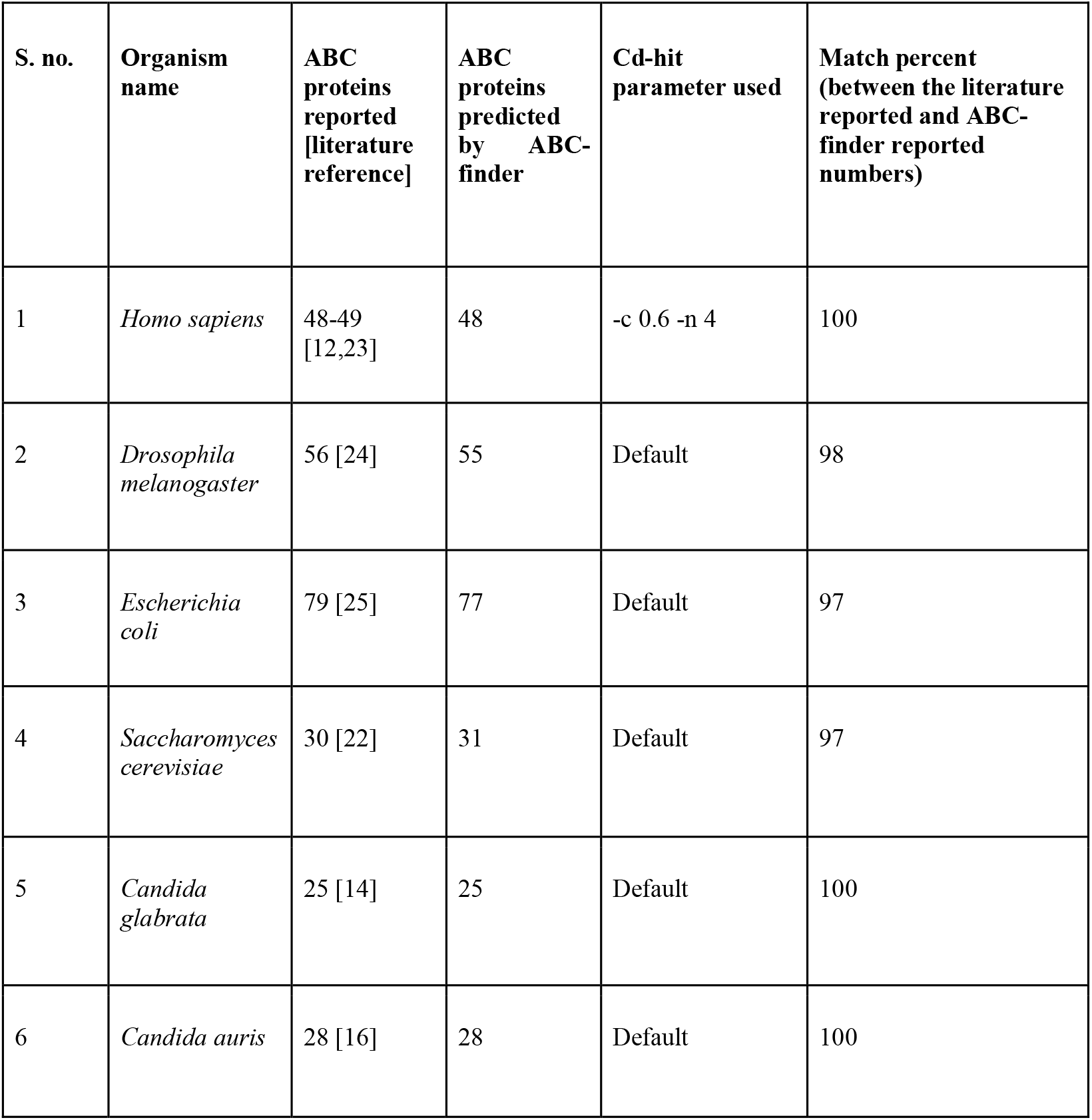
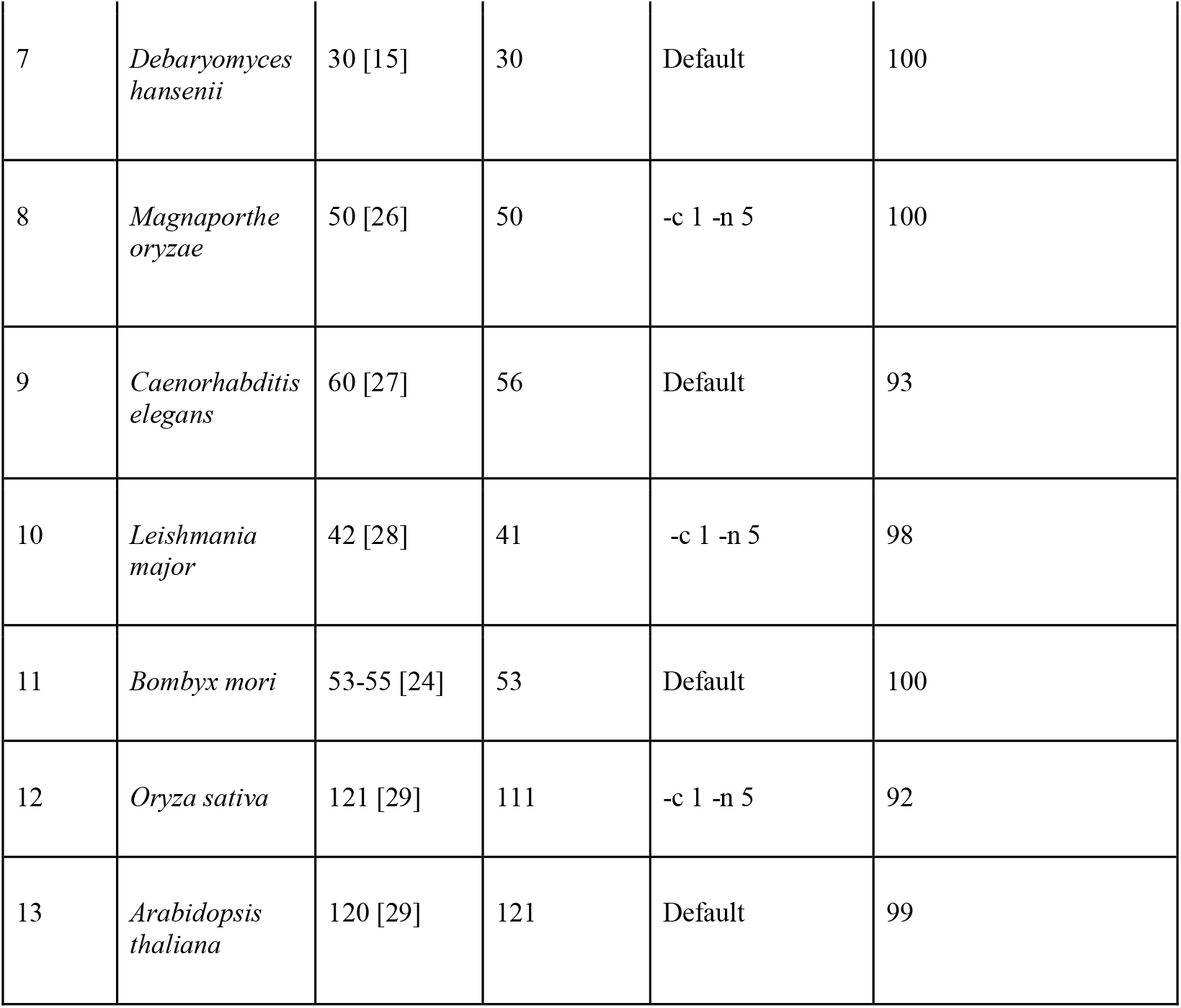
A comparison between the number of ABC proteins reported in the literature v/s predicted by ABC-finder. cd-hit parameter used: **-c =** sequence identity threshold; **-n =** word length; **Default =** (-c = 0.9; -n = 5). Result files for all the below mentioned organisms are available at https://zenodo.org/record/4603195.

Since the ABC-finder protocol uses relatively stringent parameters, the results obtained should reflect the most likely ABC candidate proteins in the user-specified proteome. To test this further, we analyzed the results for *S. cerevisiae*, most of whose ABC proteins are experimentally characterized. As per earlier studies, the organism comprises 30 ABC proteins [22], 27 of which come with some degree of experimental characterization. As summarized in **Table 2**, ABC-finder could identify all 30 of the reported ABC proteins. In addition, it also identifies one uncharacterized protein YKR104W (NP_013030.1). Further sequence and structure-function analysis will help confirm its status as an ABC protein. Since, ABC-finder utilizes TOPCONS, which combines an arbitrary number of topology predictions into one consensus prediction [20] and combines it with the results of HMM-based NBD-mapping. Hence, it presents a highly plausible domain(s) arrangement (topology) in ABC proteins **(Figure 3)**. Furthermore, the Cd-hit aided clustering, which allows the grouping of the candidate proteins according to sequence similarity, can be exploited to classify the ABC members into various subfamilies.

**Table 2:**
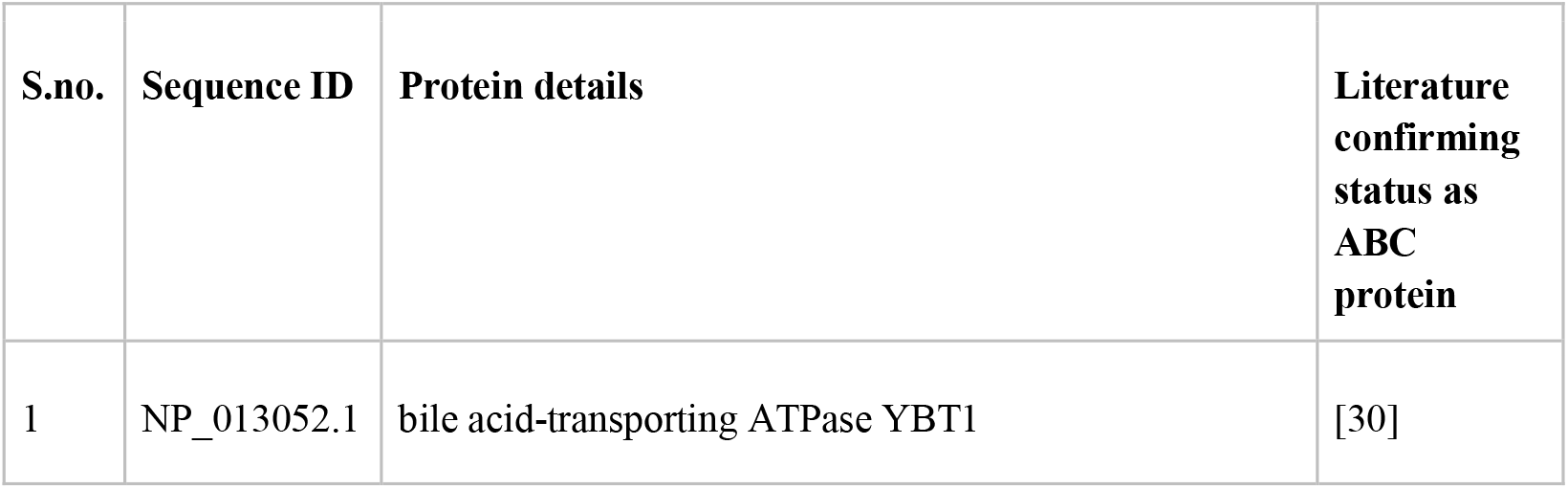
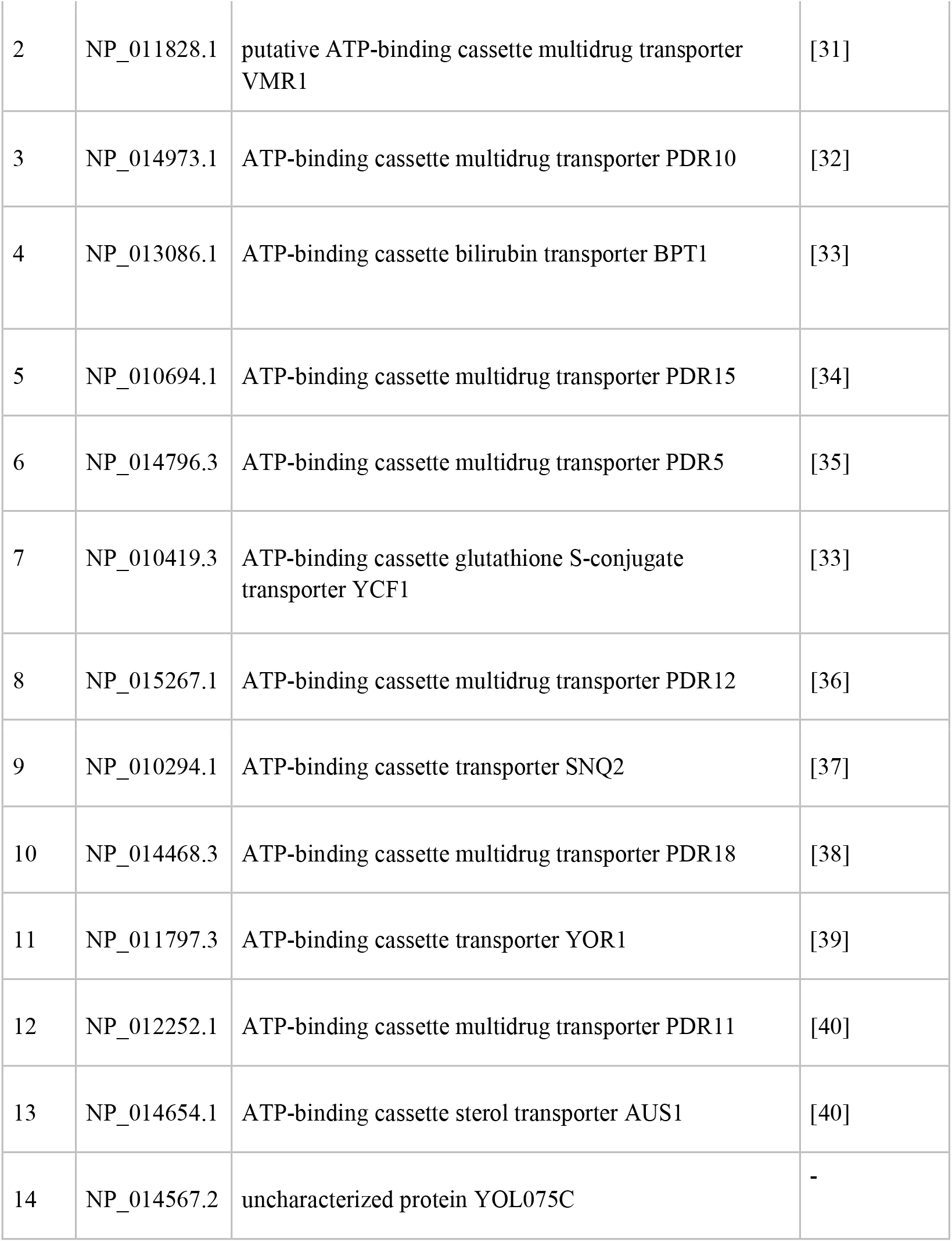
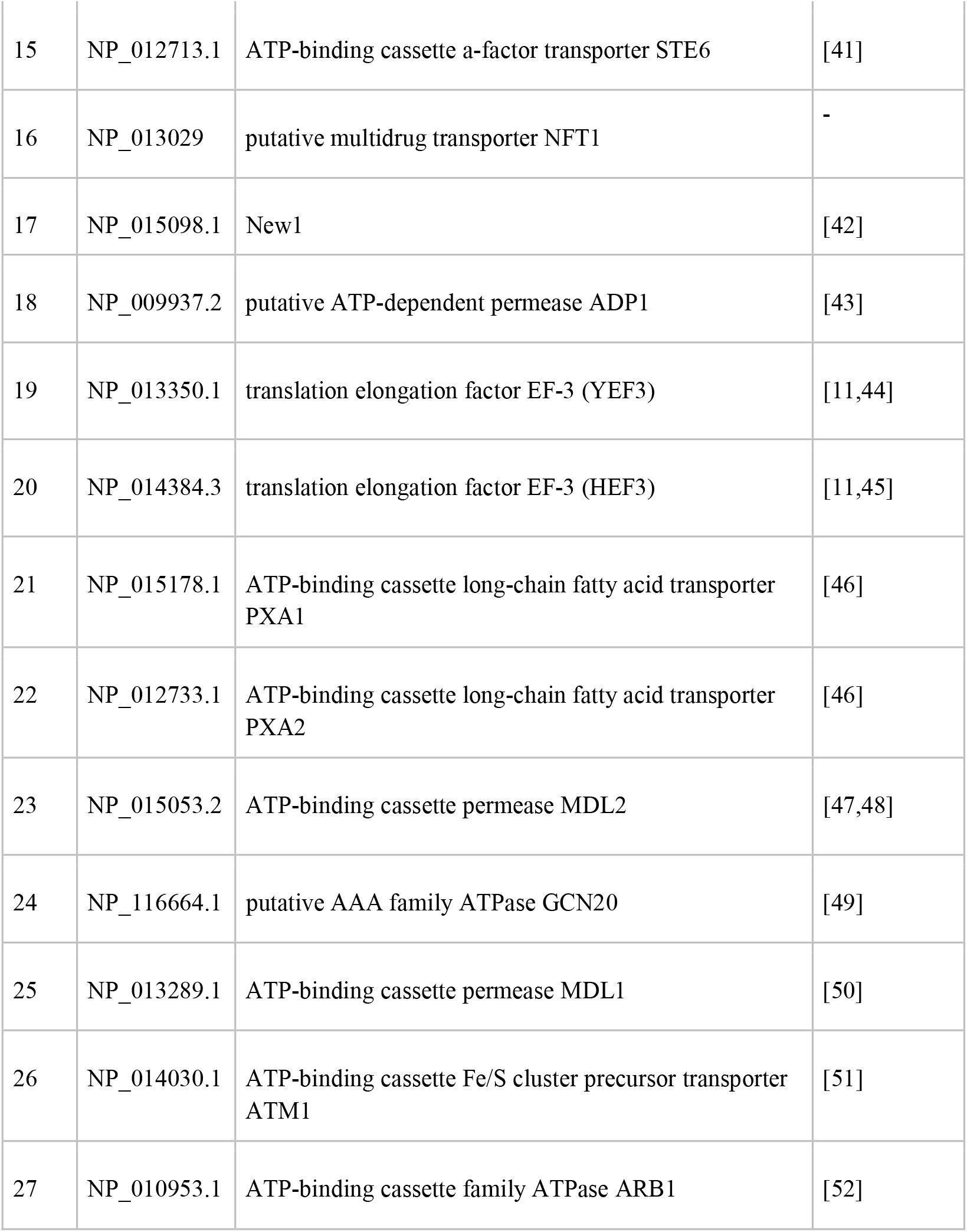
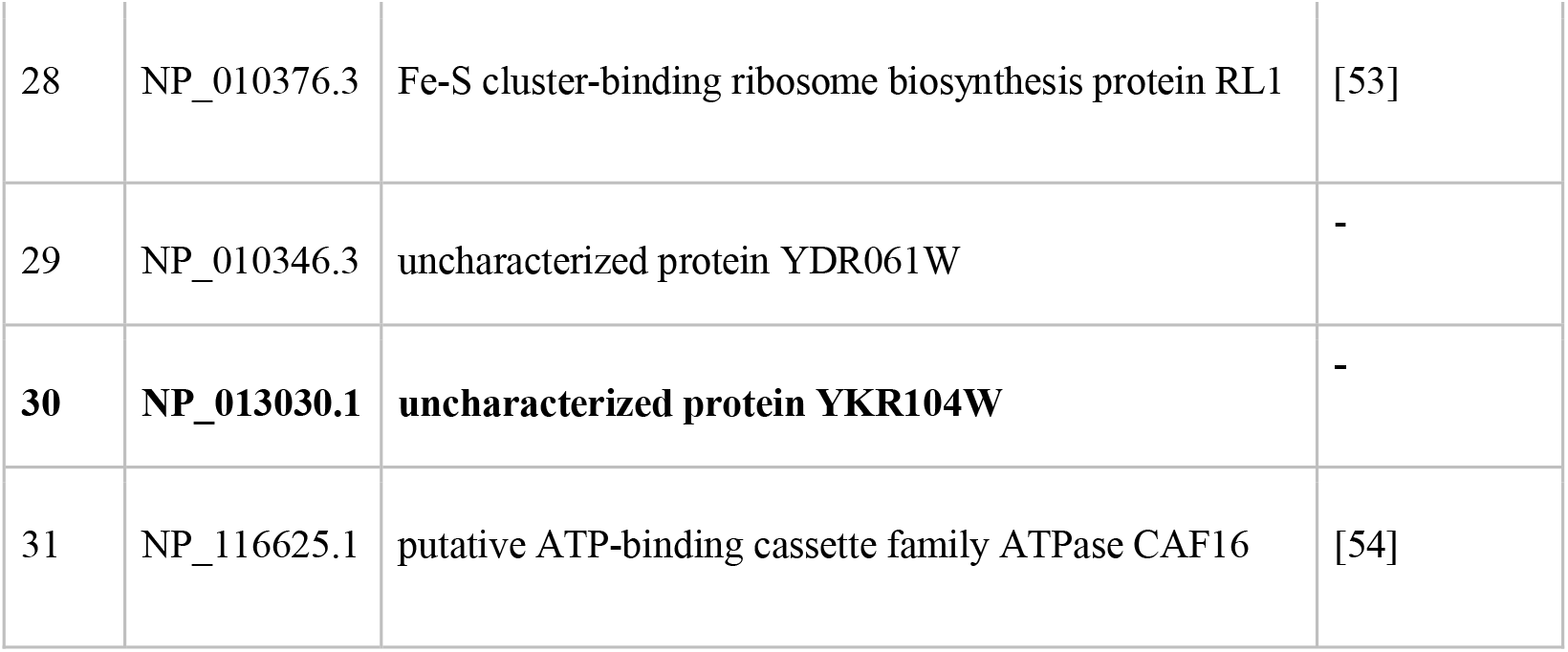
List of ABC proteins predicted by ABC-finder in the *S. cerevisiae* S288C proteome. NP_013030.1 has not been reported in previous inventories of *S. cerevisiae* ABC proteins.

Altogether, the web server caters to the ABC research community by providing a very simple tool that meets all the preliminary needs pertaining to sequence/domain analysis of the ABC proteins. Considering the need for analysis of larger datasets by some users, we have also made it feasible for the users to run ABC-finder locally. Given the numerous physiological and clinical implications of ABC proteins across the living world, the web server is an extremely useful tool that will be further improvised in the subsequent versions.

## SOFTWARE AVAILABILITY

The scripts we used to deploy the containers in our study are publicly available on GitHub and the Docker images are available on DockerHub.

### GitHub URL (project home page for source code): https://github.com/lynngroup/abcfinder

### DockerHub URL: https://hub.docker.com/r/lynngroup/abcfinder

## ACKNOWLEDGEMENTS

We are grateful to all our colleagues involved in testing the ABC-finder web server. AB acknowledges funding support from SERB (SRG/2019/000514). PV and NKM are thankful to funding agencies ICMR and Department of Biotechnology (DBT/JRF/15/AL, DBT/2016/JNU/695), Govt. of India and JNU for financial support and fellowships. The authors acknowledge the High-Performance Computational Facility (HPCF) at SCIS, JNU.

