## Supplementary information for "ABC-finder: A containerized web server for the identification and topology prediction of ABC proteins"

**Installation**

Users can install the ABC-finder web server directly from the source code or build and run it from within Docker container. Please follow the following steps:

#### **Building the Docker container**

*Clone this repository:*

$ git clone <https://github.com/lynngroup/abcfinder.git>

*Move into the repository directory:*

$ cd abcfinder

The file Dockerfile & docker-compose.yaml contains instructions for building a Docker container for ABC-finder webserver.

If the user has Docker & docker-compose installed on the computer, the container can be built. Once Docker is set up on the host computer, ABC-finder can be downloaded and installed using the following command:

$ docker pull lynngroup/abcfinder

*This will fetch the latest version with 'latest' tag.*

To run ABC-finder, use the following command:

$ docker-compose build

$ docker-compose up -d

This will initiate ABC-finder at port 8000 of the local server or localhost. The user may use another port to initiate another instance. [To manipulate Docker utilities, refer to Docker Documentation]. While the ABC-finder instance is running inside Docker container, ABC-finder User Interface (UI) can be accessed through a web browser at the following URL:

- <http://localhost:8000/> or
- [http://IP_ADDRESS_OF_HOST_COMPUTER:8000](http://ip_address_of_host_computer:8000/)

ABC-finder can now be used to upload the data using the browser.

### **Frequently Asked Questions**

1) **How do I submit a sequence to ABC-finder?**

Please follow the tutorial available on <https://youtu.be/5ckaG4cXbRw>

2) **How much time will it take to finish my job?**

It depends on the input proteome size and the number of positive hits for ABC proteins. The average time is 2.5 to 5 hours.

**Note:** Currently ABC-finder does not have “caching” feature. ABC-finder is a special case of running docker inside docker, wherein ABC-finder itself is a docker container and it’s using another docker container (TOPCONS) inside it. We will soon enable caching on ABC-finder such that faster results can be obtained in cases where someone has already run the same sequence file before.

3) **Have you tested ABC-finder on some organisms to validate the program?**

Yes, we have tested ABC-finder on 13 organisms. The result files are available at <https://zenodo.org/record/4603195>

4) **How to install ABC-finder locally**

The entire process has been described in the above-mentioned Installation section.

5) **How and whom to contact, if my job is not finished or I have any doubt?**

6) **Can you please tell components of ABC-finder**

Please check the ABC-finder related section at

<http://abc-finder.osdd.jnu.ac.in/app/related>

7) **How do I get notified about the status of my job on ABC-finder?**

If you mention your email under the RUN section of ABC-finder, you will receive notifications at the start of the job. Further, results are sent in the form of a zipped folder via email once the job is completed.

8) **Please provide your docker hub link for docker images**

The link is <https://hub.docker.com/r/lynngroup/abcfinder/tags> (we are using 4 docker images tags)

1. lynngroup/abcfinder:db
2. lynngroup/abcfinder:redis
3. lynngroup/abcfinder:web
4. lynngroup/abcfinder:celery

9) **Where to find the source code of ABC-finder?**

<https://github.com/lynngroup/abcfinder>

10) **What is the minimum hardware requirement for trying out ABC-finder locally?**

*Minimum*

- Linux 64 bits (Ubuntu/centos7).
- Any CPU (Intel i3/i5/ i7/ or Ryzen 3/5/7, quad-core→ recommended)
- 4 GB RAM, 120 GB HDD free space.

*Recommended*

- Linux 64 bits (Ubuntu/centos7).
- CPU quad-core or hexa-core Intel i7/Intel i9/ or **Ryzen** 5/7**.**
- 8 GB RAM, 130 GB HDD free space. (ABC-finder requires ~118 GB	of	TOPCONS database to be downloaded locally)

**Note:** ABC-finder uses CPU power and does not require GPU, you can easily change how many parallel jobs you want to run and what percentage of CPU you want to use. Currently, ABC finder uses max 60% of CPU. If you want to modify these settings, please send your email to **

**Important Links**

1. ABC-finder is accessible at [http://abc-finder.osdd.jnu.ac.in](http://abc-finder.osdd.jnu.ac.in/)
2. Project home page for source code: <https://github.com/lynngroup/abcfinder>
3. Docker Hub: <https://hub.docker.com/r/lynngroup/abcfinder>
